# Development of the gut microbiota during early life in premature and term infants

**DOI:** 10.1101/2022.07.24.501289

**Authors:** Kathleen Sim, Elizabeth Powell, Emma Cornwell, J Simon Kroll, Alexander G Shaw

## Abstract

**Background:** The gastrointestinal (GI) microbiota has been linked to health consequences throughout life, from early life illnesses (e.g. sepsis and necrotising enterocolitis) to lifelong chronic conditions such as obesity and inflammatory bowel disease. It has also been observed that events in early life can lead to shifts in the microbiota, with some of these changes having been documented to persist into adulthood.

A particularly extreme example of a divergent early GI microbiota occurs in premature neonates, who display a very different GI community to term infants. Certain characteristic patterns have been associated with negative health outcomes during the neonatal period, and these patterns may prove to have continual damaging effects if not resolved.

**Results:** In this study we contrast a set of premature infants with a paired set of term infants (n=37 pairs) at six weeks of life and at two years. In the samples taken at six weeks we find microbial communities differing in both diversity and specific bacterial groups between the two infant cohorts. We identify clinical factors associated with over-abundance of potentially pathogenic organisms (e.g. Enterobacteriaceae) and reduced abundances of some beneficial organisms (e.g. *Bifidobacterium*).

We contrast these findings with samples taken at two years of age, which show that despite a very different initial microbiota, the two infant groups converge to a similar, more adult-like state. We identify clinical factors, including both prematurity and delivery method, that remain associated with components of the microbiota. Both clinical factors and microbial characteristics are compared to the occurrence of childhood wheeze and eczema, revealing associations between components of the GI microbiota and the development of these allergic conditions.

**Conclusions:** The faecal microbiota differs greatly between infants born at term and those born prematurely during early life, yet it converges over time. Despite this, early clinical factors remain significantly associated with the abundance of some bacterial groups at two years of age. Given the associations made between health conditions and the microbiota, factors that alter the makeup of the gut microbiota, and potentially its trajectory through life, could have important lifelong consequences.

## Background

The gut microbiota is a vast reservoir of bacteria living within a human host with interactions both beneficial and harmful. The community plays an important role in the digestion of food, the development of the immune system and in the maturation of the gut itself [1-3]. By the point of birth, the gut has been seeded with an array of organisms that colonize and grow, only to be replaced by a succession of species as the environment changes[4, 5]. Great variety is seen in the diversity and abundance of organisms, and the patterns present have been associated with health conditions throughout life [6]. The causes of variation within the community are therefore of great interest in trying to predict- and potentially enhance-future health, and it has been indicated that factors in early life such ais mays mode of delivery can still be reflected and observed in the patterns of the gut microbiota in adults [7].

We may therefore be concerned by the particularly extreme case of gut microbiota alteration that occurs as a result of the net effects of being born premature. These infants are found to have a drastically altered gut microbiota when compared to infants born at term [8, 9]. Prior research has demonstrated that particular community patterns are associated with conditions such as NEC and sepsis in early life, as well as conditions such as IBD and obesity in later life [4, 10-13]. The trajectory of the infant gut microbiota is therefore of great interest, with potential life-long consequences.

A particular area of interest for the infant microbiota is the potential relationship with allergic disease. The rapid rise in allergic disease including asthma and eczema may have an origin in the changing exposure to the environmental microbiota in recent years [14]. Environmental biodiversity and the individual’s early microbiota may affect the development of immune tolerance [14], and a critical window when the microbiota may exert its influence has been supported by both animal models [15] and cohort studies [16, 17]. Reduced diversity of the faecal microbiota has been associated with allergic sensitization [18], rhinitis [18] and later asthma [17] though not uniformly in all studies. Specific bacterial taxa present early in the faecal microbiota such as clostridium have been associated with an asthma diagnosis [19] and allergic sensitization [20], and conversely reduction in other taxa such as lachnospira have been associated with asthma; with this association confirmed in a mouse model [16].

In this study we sought to determine if the gut microbiota of term and premature infant cohorts converge by two years of age, and to study whether microbial signatures that differ between the cohorts are associated with wheezing and eczema.

## Methods

### Study Population and sample collection

#### Preterm cohort

Premature infants (< 32 weeks gestation) who were admitted to an Imperial College Healthcare NHS Trust neonatal intensive care unit (NICU) (St Mary’s Hospital, Queen Charlotte’s and Chelsea Hospital) between January 2011 and December 2012 were recruited to the Neonatal Microbiota (NeoM) Study. The study was approved by the West London Research Ethics Committee Two, United Kingdom (reference number: 10/H0711/39). Parents gave written informed consent for their infant to join the study. Detailed daily clinical data was collected during the duration of the admission. We aimed to collect every faecal sample produced by each participant from recruitment until discharge. Nursing staff collected the samples from the nappies using a sterile spatula and placed them into a sterile DNAase-, RNAase-free microcentrifuge tube. Samples were stored at -20°C within two hours of collection and transferred to a -80°C freezer within five days, where they remained until DNA extraction.

Parents were re-approached when the participants were between the ages of two and four years to join the follow-up study (NeoM2) (approved by the London – Chelsea Research Ethics Committee - reference number: 13/LO/0693). Parents who consented to their child participating in the study completed a general health questionnaire for their child which contained questions specifically relating to allergic disease and wheezing episodes. A faecal sample from the child was also collected by the parents using a sterile scoop and placed into a sterile container which was then posted to the research laboratory within 24 hours. Faecal samples were immediately stored at -80oC on receipt, where they remained until DNA extraction.

#### Term cohort

Expectant parents were approached in the antenatal clinics of St Mary’s hospital and asked for their assent to be approached when their baby was born. Subsequently, if the babies met the inclusion criteria (≥37 weeks gestation with no airway abnormalities and born at St Mary’s hospital), parents were re-visited on the postnatal ward or at home and their written informed consent was asked for their baby’s participation in the Development Of the Respiratory Microbiota (DORMIce) study. A smaller proportion was recruited directly on the postnatal ward. The study was approved by the London Riverside Research Ethics Committee, United Kingdom (Reference number: 13SM0646).

The study involved regular visits to the infants in the community at 6 weeks, 6, 9, 12, 18 and 24 months, and in a subset at 3 and 4.5 months. Faecal samples were collected at birth and at each timepoint by the parents from the nappy using a sterile scoop and placed into a sterile container. This was transported by the researcher to the laboratory or alternatively posted by the parents and subsequently frozen at -80°C.

For each cohort, a sample closest to 6 weeks after birth and one taken at 24 months were chosen for analysis. Clinical data describing the infant’s feeds, living environment and health history was collected at each of the timepoints either by reference to clinical notes or through a questionnaire. The assembled data for the term and premature infant cohorts is summarised in Table 1.

**Table 1.** Demographics of the DORMICe (term infants) and NeoM (premature infants) cohorts at birth, six weeks and two years of chronological age. For continuous factors median and interquartile range are shown. Categorical factors are expressed in absolute value and percentage of the cohort in brackets.

|  | **DORMICe** | **NeoM** |
| --- | --- | --- |
| ***Birth Demographics*** |  |  |
| Female | 19 (51.4) | 14 (37.8) |
| Median birth weight (gram) | 3510 (3130, 3690) | 960 (765, 1325) |
| Mean gestation at birth (weeks +days) | 40+2 (39+4 to 41+2) | 28+1 (25+4 to 29+6) |
| **Ethnicity** |  |  |
| Black | 2 (5.4) | 4 (10.8) |
| White | 20 (54.1) | 18 (48.4) |
| Asian | 4 (10.8) | 9 (24.3) |
| Mixed race | 6 (16.2) | 3 (8.1) |
| Other | 5 (13.5) | 3 (8.1) |
| **Mode of Delivery** |  |  |
| Caesarean section | 8 (21.6) | 8 (21.6) |
| Vaginal | 29 (78.4) | 29 (78.4) |
| ***Six Week Demographics*** |  |  |
| **Feeding at time of Sample** |  |  |
| Breast milk | 26 (70.3) | 26 (70.3) |
| Formula | 10 (27.0) | 10 (27.0) |
| Mixed | 1 (2.7) | 1 (2.7) |
| **Number of complete months of breast milk feeds** | 1 (1, 1) | 1 (0, 1) |
| **Number of courses of antibiotics** | 0 (0, 0) | 1 (1, 2) |
| ***Two Years Demographics*** |  |  |
| **Infant age at sampling (months)** | 24 (23.9, 24.2) | 35.7 (31.1, 39.1) |
| **Feeding at time of Sample** |  |  |
| Weaned | 37 (37, 37) | 37 (37, 37) |
| **Number of complete months of breast milk feeds** | 11 (7,17) | 6 (3, 9) |
| **Age had non-breast milk feeds (months)** | 0 (0, 4) | 3 (0, 6) |
| **Number of courses of antibiotics since six weeks** | 1 (0, 2) | 3 (0, 4) |
| **Antibiotics use in the last month (yes)** | 2 (5.4) | 2 (5.4) |
| **Pet in the household (yes)** | 7 (18.9) | 10 (27.0) |
| **Other child in the household (yes)** | 15 (40.5) | 21 (56.8) |
| **Smoker in the household (yes)** | 5 (13.5) | 2 (5.4) |
| **Child suffered from one or more wheezing episodes (yes, parent reported)** | 18 (48.6) | 13 (35.1) |
| **Child suffers from eczema (yes, parent reported)** | 24 (64.9) | 7 (18.9) |

#### Matching Criteria

Thirty-seven infants from the NeoM study were matched to 37 infants from the DORMICe study by mode of delivery and the type of feeds (breast milk/formula/mixed feeds) that the infant was on at the time of collection of the 6 week faecal sample. Where possible, infants were also matched on maternal intrapartum antibiotic use and neonatal antibiotic use at birth, followed by ethnicity.

#### Bacterial DNA extraction

DNA was extracted from the faecal samples (200mg) using the FastDNA SPIN Kit for Soil, following the manufacturer’s protocol (inclusive of bead-beating homogenisation steps) except that the final elution step was into Tris (10mM) low-ethylenediaminetetraacetic acid (0.1mM) buffer.

#### Amplification and pyrosequencing of the V3-V5 regions of the bacterial *16S rRNA* gene

Primers containing a unique 12-bp Golay barcode [21, 22] were used to amplify the V3-V5 region of the bacterial *16S rRNA* gene from each DNA sample. Amplicons were produced by polymerase chain reaction (PCR) as described previously [23]. The resulting pooled replicate amplicons were purified and three pyrosequencing runs were conducted on a 454 Life Sciences GS FLX (Roche) machine in accordance to the Roche Amplicon Lib-L protocol. Negative extraction controls were included to identify contamination. An average of 1650 sequencing reads was obtained per sample.

#### Data Processing

Shotgun processed data were analysed using the Quantitative Insights Into Microbial Ecology (QIIME) version 1.9.0 package [24], following the recommended pipeline for the combination of multiple 454 FLX datasets. Denoising was performed using denoise_wrapper.py, and the datasets integrated. Chimera removal was performed with ChimeraSlayer [25]. Sequences were clustered at 97% sequence identity using the UCLUST algorithm into OTUs [26] and aligned to the SILVA rRNA database version 119 [27]. Rarefaction to 664 reads per sample was performed, removing heterogeneity of sequencing reads per sample.

#### Statistics

Statistical analyses were performed with the R statistical package [28]. Alpha and Beta diversity measures were calculated using QIIME [24]. Beta diversity distances between sample groups were compared using the Mann-Whitney U test (testing distances within groups to between groups) and the Wilcoxon signed-rank test (testing matched sets of distances at different time points). Alpha diversity was compared using the Wilcoxon signed-rank test. Canonical ordination analysis (CCA) was performed in R with the vegan statistical package [29].

Differentially abundant OTUs/phyla at six weeks and two years of age were detected using general linear models with a negative binomial distribution, with weeks gestation at birth used as the explanatory variable and day of sampling added as a confounding factor. Summary tables for OTUs (Tables 1 and 5) only contain OTUs found to be significantly different, whilst information for all phyla is presented (Tables 2 and 6). Multiple hypothesis corrections (MHC) were made with the Bonferroni correction.

**Table 2.**
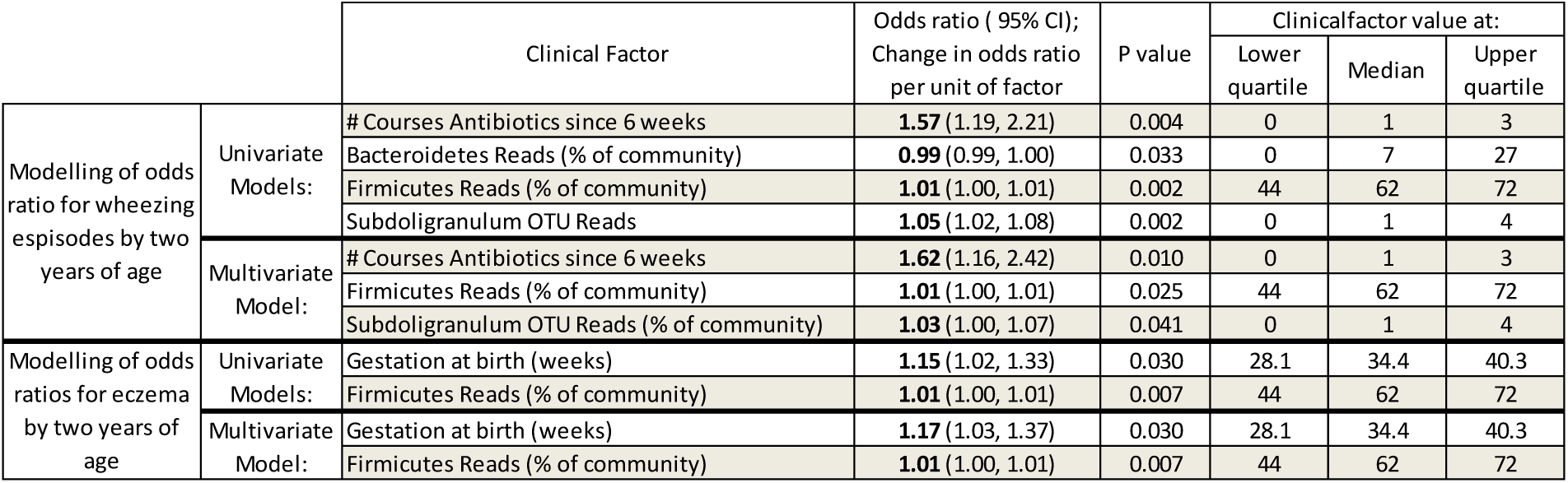
Results of models relating microbial data and clinical factors to the odds of wheezing episodes and eczema at two years of age. Univariate models included the investigated variable, infant gestational ages and day of sampling and factors have been reported if significant at the 20% level after MHC (OTUs, phyla and clinical factors considered separate groups). A multivariate model was used to identify dominant factors, with iterative removal of the least influential factors until only significant factors remained. Odds ratios and 95% confidence intervals have been calculated for each model; values provided indicate the proportional change in odds per unit of the variable.

Missing clinical data was first imputed using the MICE package through predictive mean matching [30]. Associations between clinical factors and OTUs/phyla were tested for using generalised linear models (GLM) with the clinical factor and as the explanatory variable and weeks gestation at birth and day of sampling added as confounding factors. MHCs were made with the Bonferroni correction, and factors that were significant at 20% were carried forward. Where multiple factors were found to be significantly associated with OTUs/phyla, multivariate models were used. Models were then refined by sequential removal of the least significant factors until only factors significant at 5% remained. OTUs/phyla that are significantly associated with clinical factors through this method are documented in Tables 4, 5, 8 and 9. Gestation at birth and day of sampling were retained in each model, with gestation at birth being documented if it significantly influenced the model.

Associations between eczema/wheeze occurrence and clinical factors and OTUs/phyla was tested for using logistic regression models, again including gestation at birth and day of sampling added as confounding factors. Corrections were performed as for the GLM models, and univariate and multivariate models were processed in the same manner, although gestation at birth was only retained in the final multivariate model if found to provide significant improvement. ROC curves and area under the curves (UAC) were calculated using R. Validation of the models was performed using sets of 30 term and 30 premature infants drawn randomly from a pool of 64 premature and 35 term infants whose data was not used elsewhere in this manuscript. Infants were drawn from the pool 1,000 times and the median ROC curve and inter-quartile range calculated.

## Results

Following exclusion of samples with insufficient sequencing reads from consideration, 148 samples remained, comprising 37 infant pairs (one term and one premature infant) each with sequencing data from samples taken at six weeks and two years. Sequencing reads assigned to the 26 most abundant OTUS (making up 95% of the total reads in the dataset) are shown segregated by term/premature birth status and by timepoint in Figure 1.

**Figure 1.**
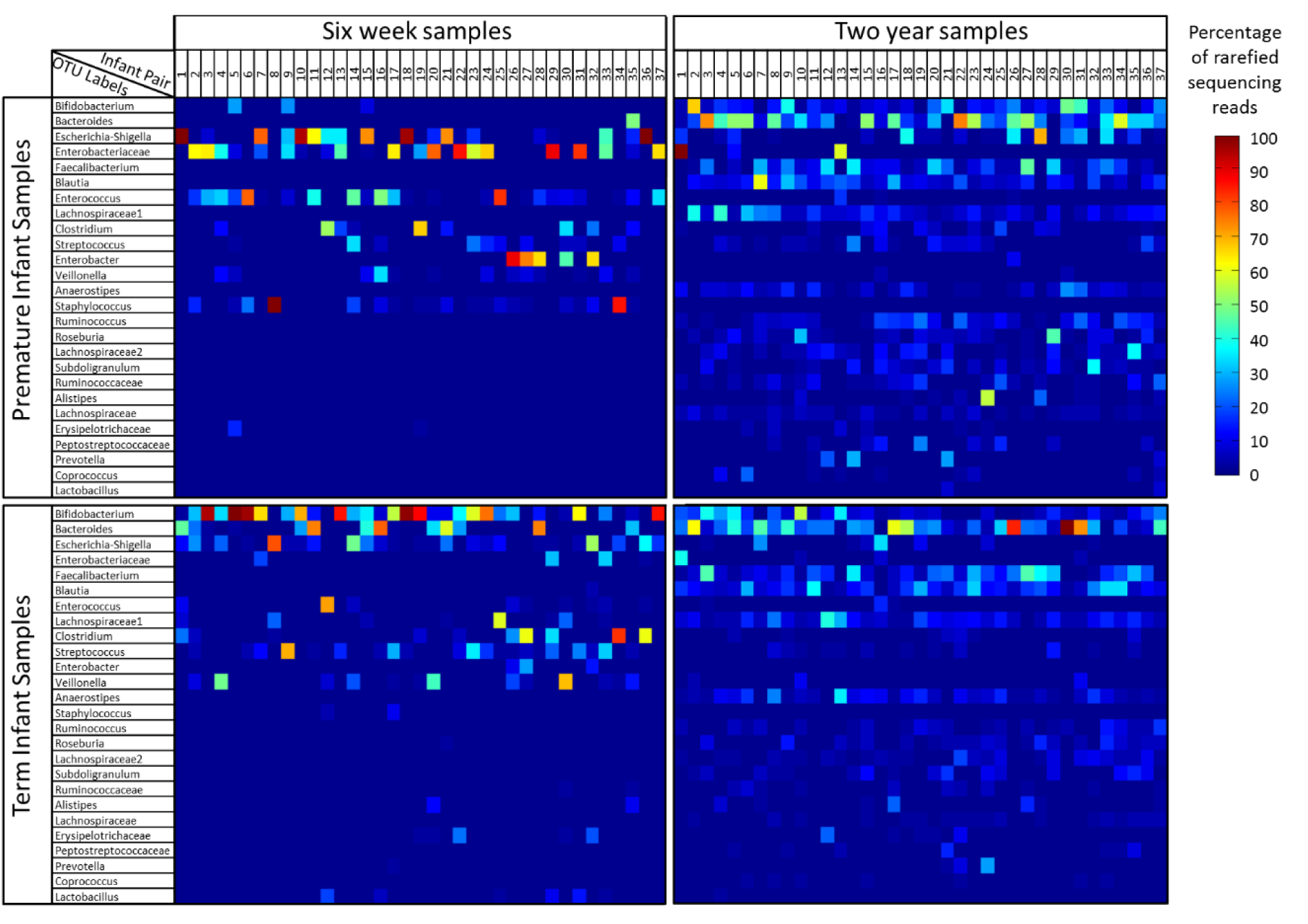
Heatmaps of rarefied sequencing data. The dataset is split into four panels according to the infant birth category (premature or term delivery) and the timepoint at which it was taken (six weeks or two years). The distribution of sequencing reads for each sample is represented by a single column in one of the four panels, with colours indicating the percentage of reads assigned to each of the OTUs shown on the y axis. Within each panel, samples are organised along with x axis by the infant pair number. Comparing columns of data vertically therefore contrasts a premature and term infant pair of samples at a single timepoint, whilst comparison of columns of data horizontally between each panel allows assessment of the changes in an infant’s faecal microbiota from six weeks to two years of age.

### The faecal microbiota of premature and term infants are significantly different at six weeks of age

To determine how similar the sample group were, distances between samples were calculated using four measures of beta diversity. These distances were plotted in principal coordinate analyses (see Figure 2) which demonstrate clear separation of the premature and term infant faecal microbial communities at six weeks by each of the four measures.

**Figure 2.**
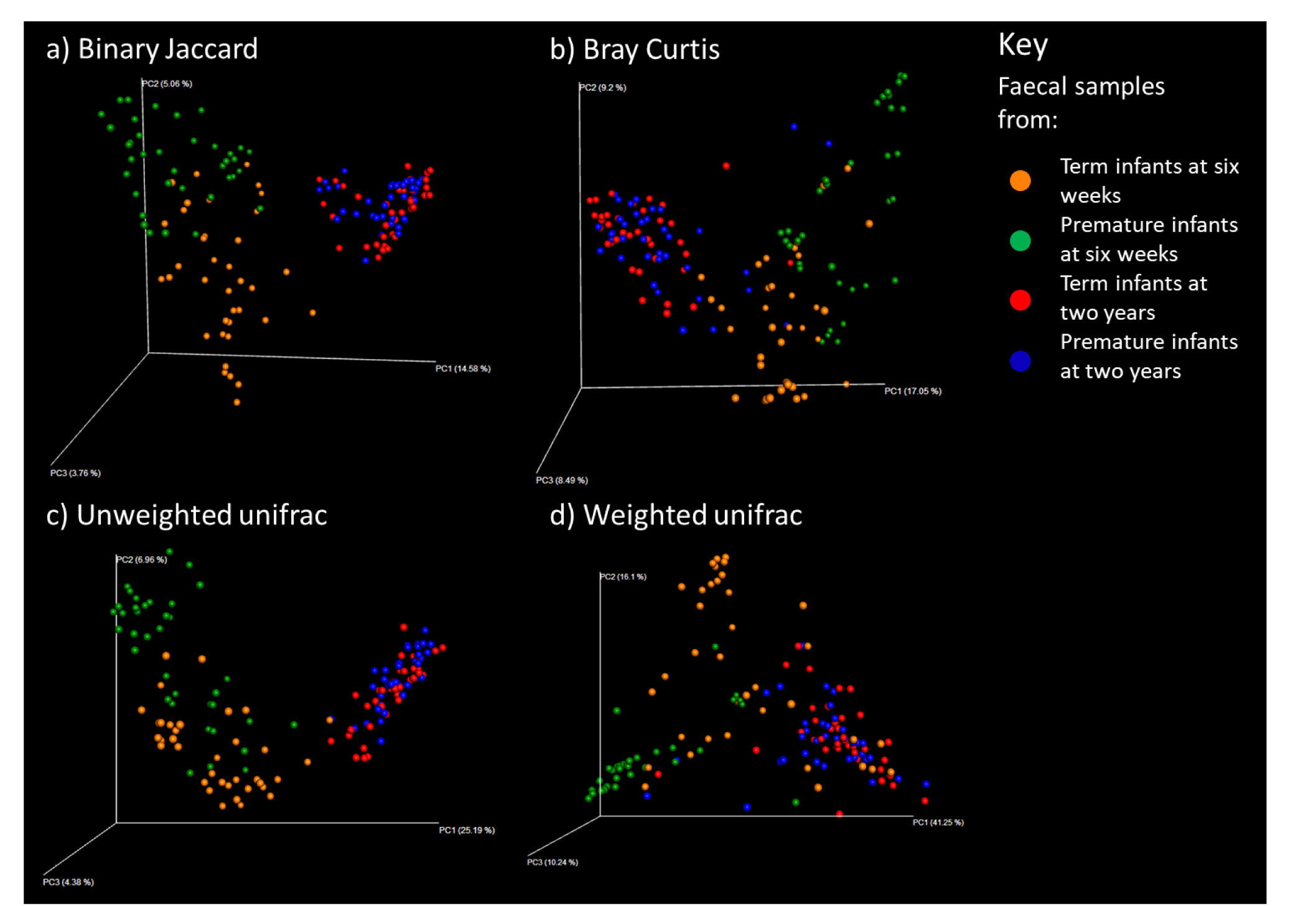
Measures of distances between samples displayed by principal coordinate analysis. Samples are colour coded according to the key.

The distances between samples within the groups were found to be significantly lower than the inter-group distances (see Figure 3), indicating that there are significant sources of dissimilarity between the microbiota of term and premature infants at six weeks of age.

**Figure 3.**
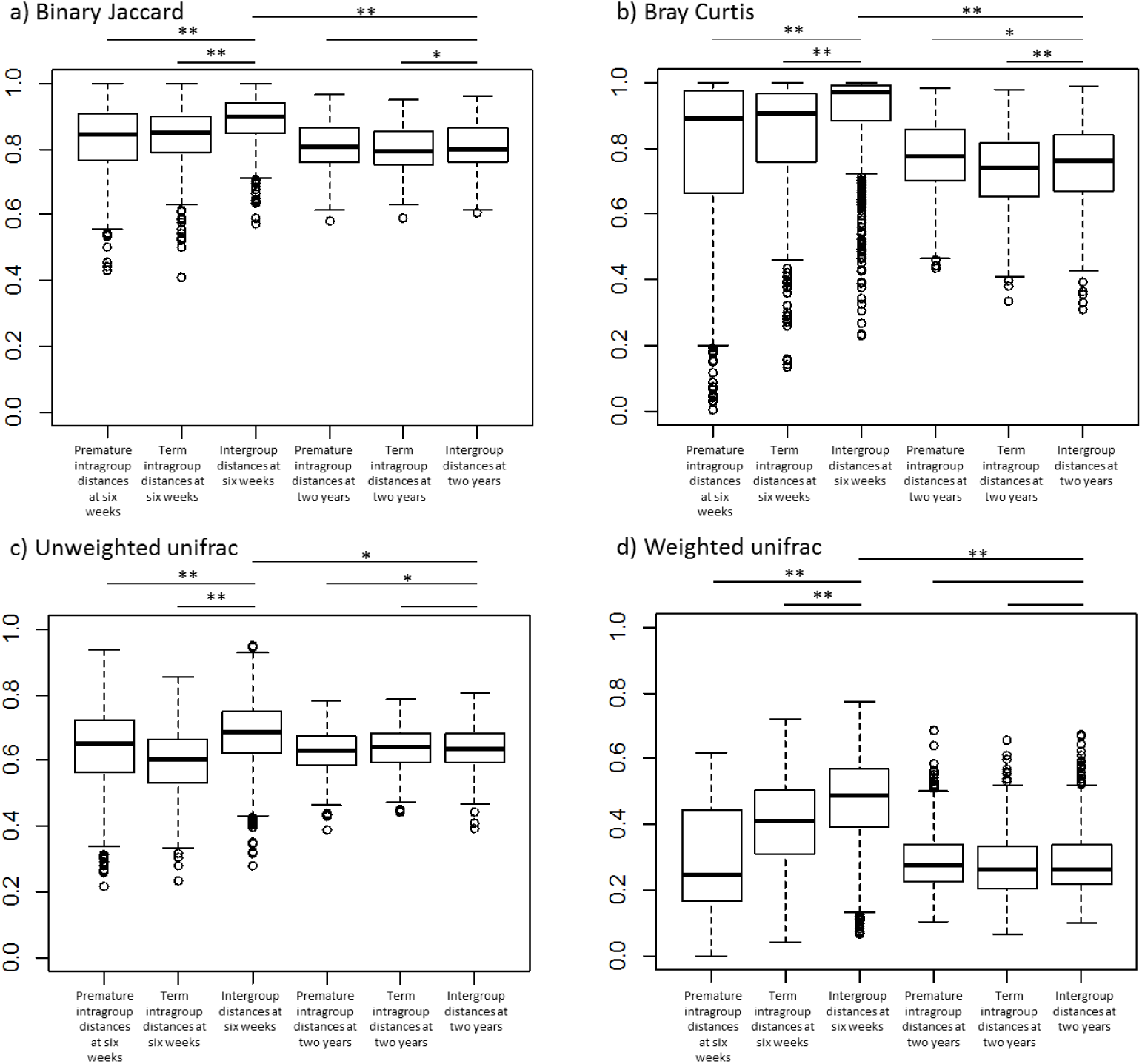
Comparisons of similarity between sample groups using four distance measures. Four sample groups have been analysed; samples from premature infants at six weeks, samples from premature infants at two years, samples from term infants at six weeks and samples from term infants at two years. Boxes indicate the 25% and 75% quartiles with the bar showing the median. Whiskers mark the quartiles ± 1.5 IQR. The x axis indicates the set of samples for which distances have been measured. Bars along the top of the charts indicate where a comparison has been made, either using a Mann Whitney U test (intragroup to intergroup distances) or a Wilcoxon signed-rank test (intragroup distances at six weeks to intragroup distances at two years) has been made. Where statistical tests found a significant difference, stars indicate the p value: * indicates 0.05 > p ≥ 0.001, ** indicates p < 0.001.

### Both diversity measures and specific OTUs differ between premature and term infant faecal microbial communities

Given the significant differences between the faecal microbial communities of premature and term infants at six weeks of age, we sought to characterise which traits of the community drove the separation.

Three diversity measures were calculated for each sample, with the results shown in Figure 4. Comparisons between premature and term infant faecal samples at six weeks indicated a significant decrease in diversity by all measures in the premature samples, reflecting a community that is both less rich and less even.

**Figure 4.**
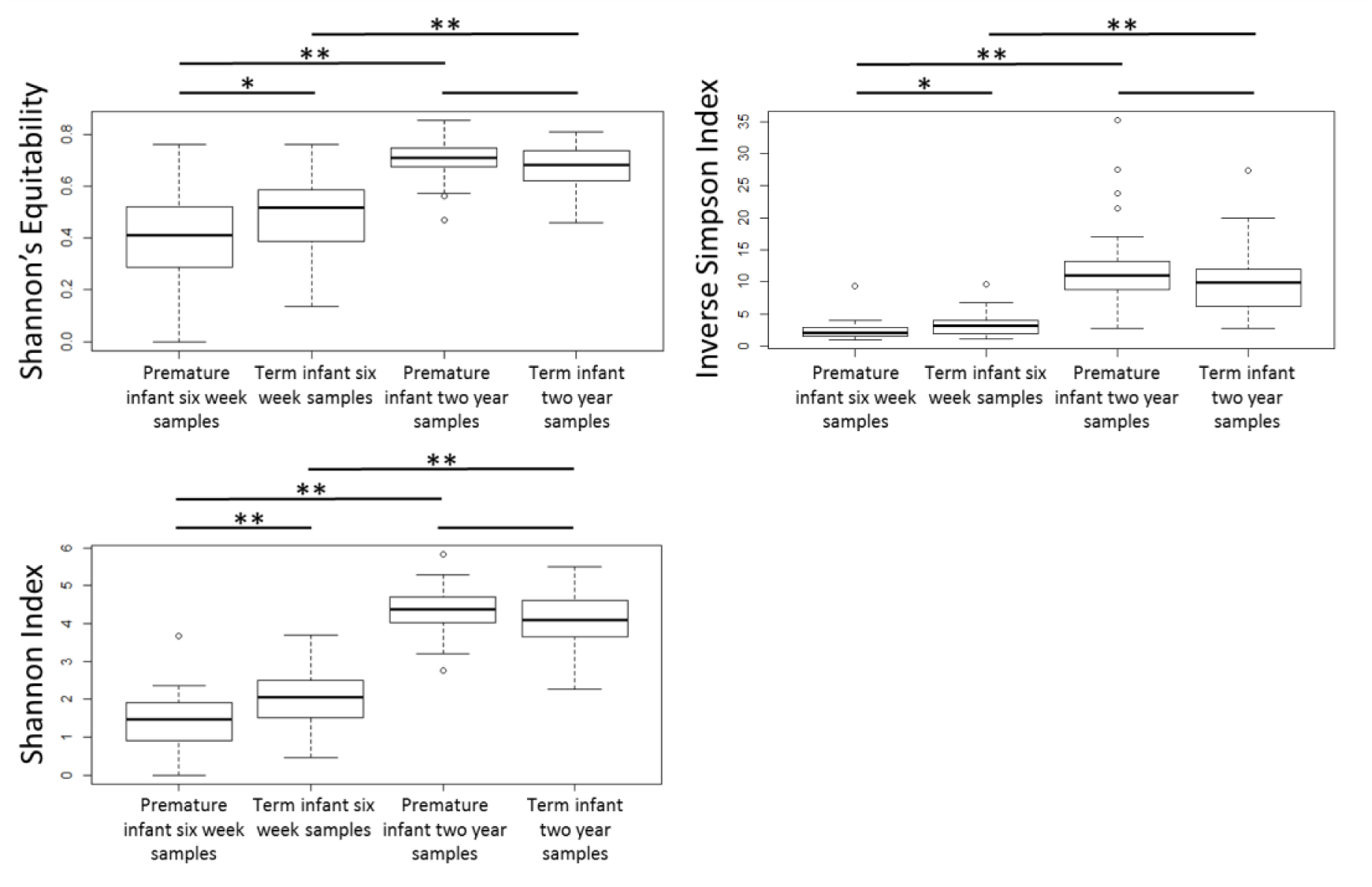
Diversity of the faecal microbial community in four groups of samples as measured by the Shannon’s Equitability, the inverse Simpson Index and the Shannon Index. Boxes indicate the 25% and 75% quartiles with the bar showing the median. Whiskers mark the quartiles ± 1.5 IQR. The diversity of sample groups was compared using the Wilcoxon signed-rank test, Where significant difference in diversity measures were found, stars indicate the p value: * indicates 0.05 > p => 0.001, ** indicates p < 0.001.

A Canonical Correspondence Analysis (CCA) of the OTUs indicates that individual bacterial groups are also driving separation between the term and premature infants (Figure 5).

**Figure 5.**
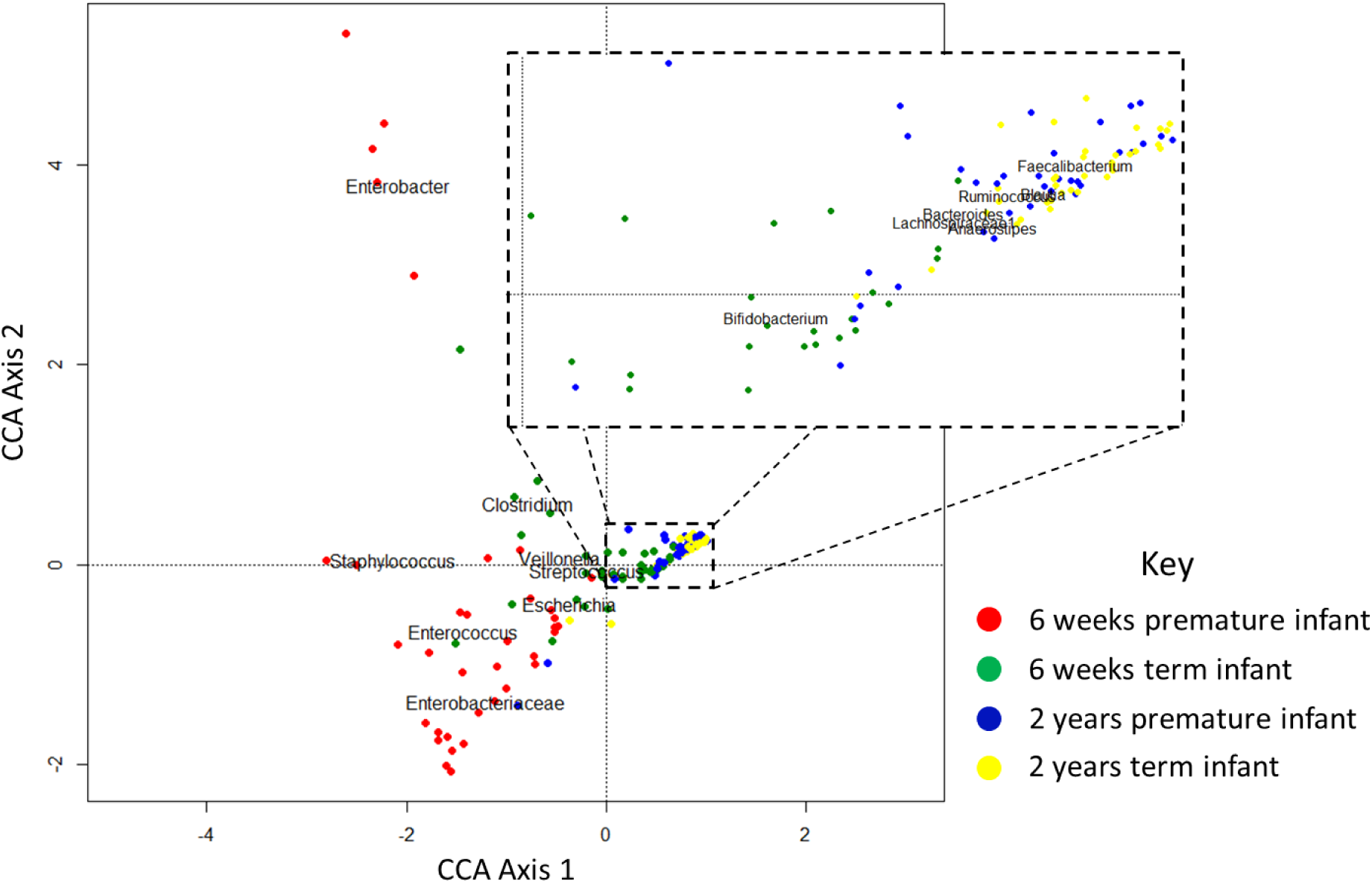
CCA showing separation of samples based on OTUs. Faecal samples taken at six weeks from premature infants are distinguished from term infants by higher abundances of Staphylococcus, Enterococcus and Enterobacteriaceae species. The microbiota of term infants feature greater abundance of Clostridium and Bifidobacterium species. By two years, the communities of both cohorts converge, with domination by genera such as Faecalibacterium, Blautia and Bacteroides.

To quantify the variation of specific OTUs between term and premature infants, we sought associations between gestational age and i) the 26 most abundant OTUs (see Supplementary Table 1) and ii) phyla (Supplementary Table 2) using GLMs. *Bifidobacterium, Bacteroides* and *Lachnospiraceae* were significantly positively associated with gestational age, whilst Enterobacteriaceae were significantly negatively associated. These findings are reflected at the phyla level, with gestational age being significantly positively associated with Bacteroidetes and Actinobacteria and negatively associated with Proteobacteria. Model parameters were used to predict the average community structure for an average term (28.1 weeks gestation) and an average premature infant (40.3 weeks gestation) at six weeks of age (See Figure 6a and 6b).

**Figure 6.**
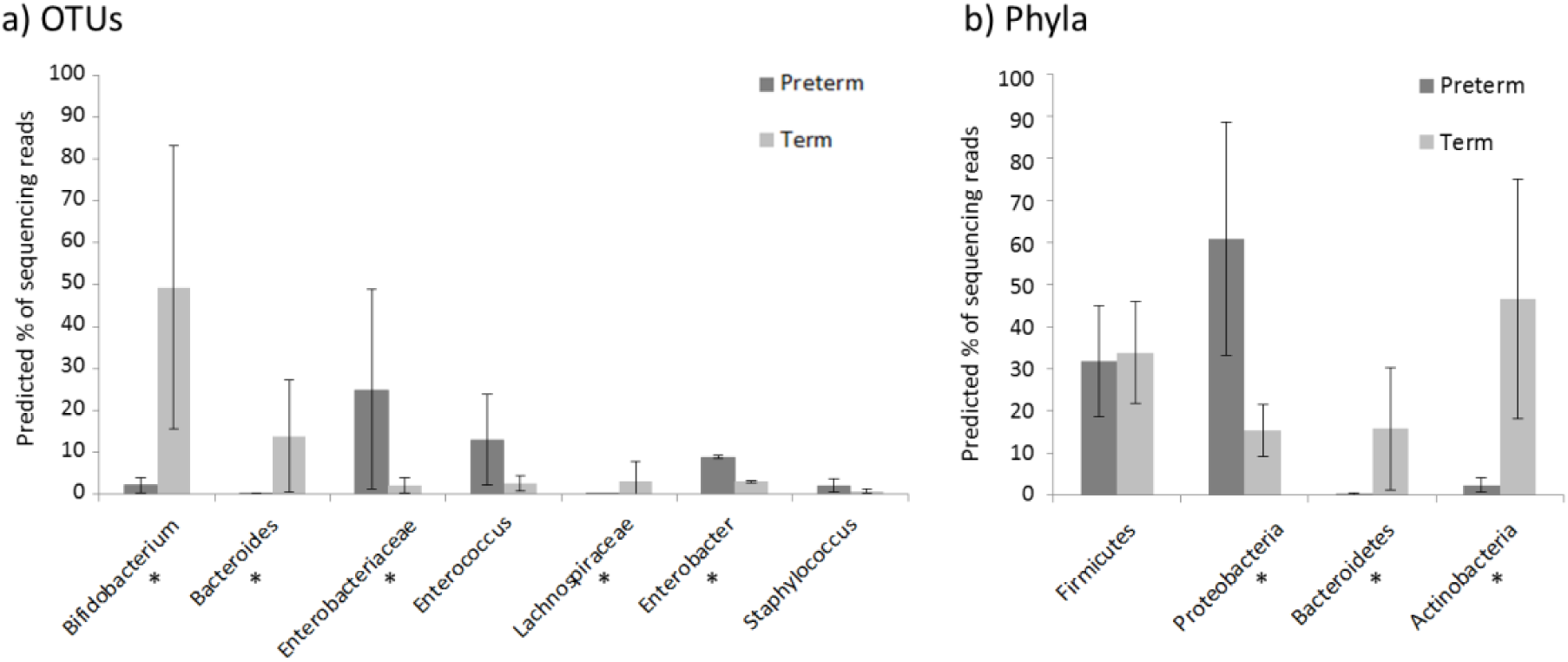
a) Bacterial OTUs that significantly differ between term and premature infants and b) Bacterial phyla in term and premature infants faecal samples. The predicted percentage of sequencing reads (of the total in the bacterial community) for each OTU at six weeks of life have been calculated using the average gestational age of premature and term infants in this dataset (27.76 weeks and 40.35 weeks respectively). Predicted values are indicated by bars, with whiskers indicating the 95 % confidence interval. In a) OTUs are displayed if significantly associated with age in GLMs, whilst in b) all phyla are shown. Stars indicate that the association with age remained significant after MHC.

### Differences in the bacterial communities can be associated with specific clinical factors that differ between being born prematurely or at term

Having shown that the bacterial community is different in term compared to premature infants, we sought to determine if the differences were primarily due to prematurity itself or were more closely associated with clinical factors that differ between infants born at term and those born prematurely.

OTUs were tested for associations with the following set of clinical factors: Feeding method at time of sampling (breast milk, formula or mixed), number of complete months of breast feeding, number of courses of antibiotics, birth method, gender, birth weight and gestation at birth (as shown in Table 1). Factors found to be significant after MHC are shown in Figure 7a; where multiple factors were found to be associated with a single OTU, a multivariate model was used to identify dominant factors, with iterative removal of the least influential factors until only significant factors remained (see Supplementary Table 3). The same process was also performed with regard to phyla rather than OTUs (see Figure 7b and Supplementary Table 4).

**Figure 7.**
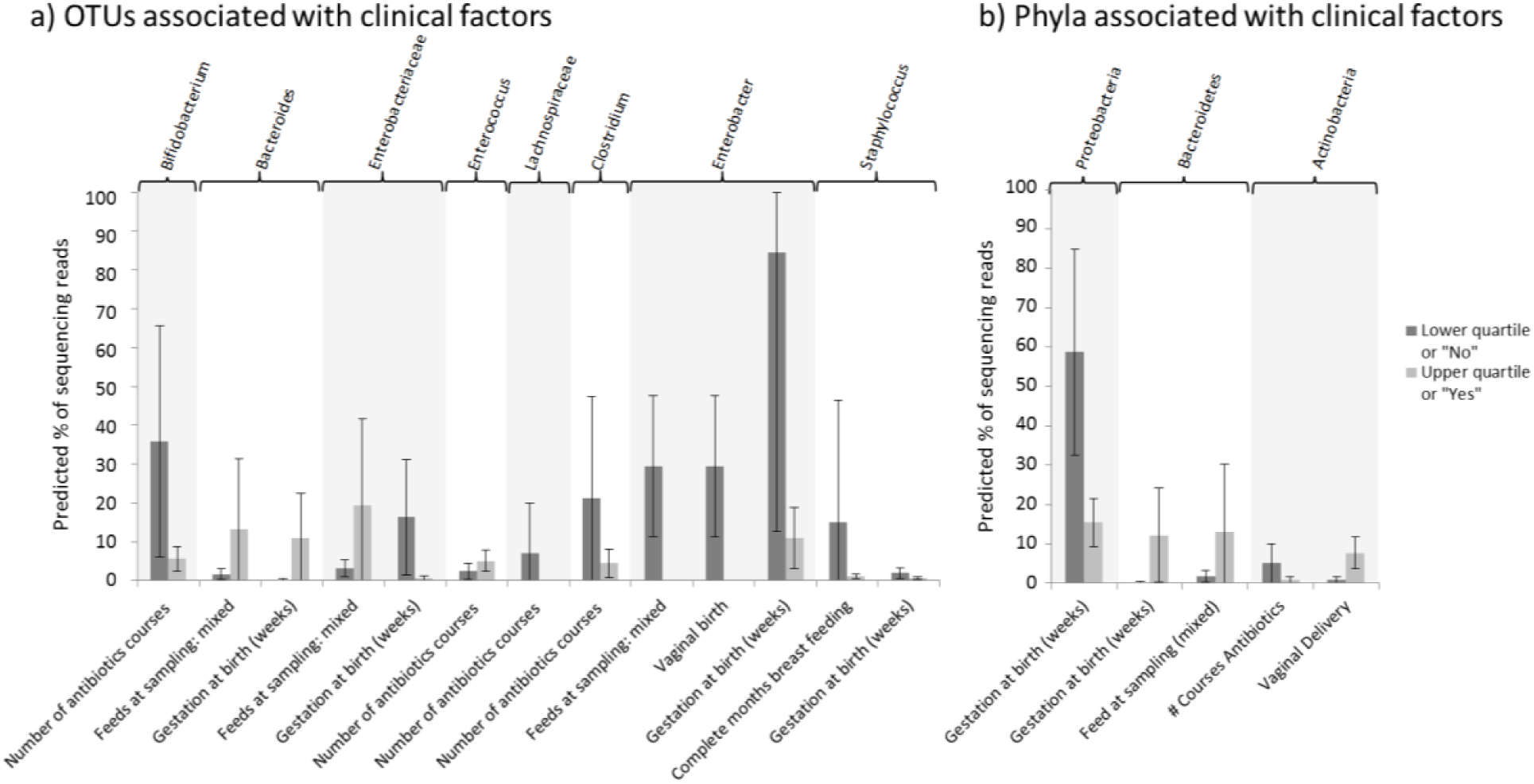
a) OTUs and their significantly associated clinical factors at six weeks of age. b) Phyla and their associated clinical factors at six weeks of age. Univariate and multivariate models were built where significant associations were found between clinical factors and OTU/phyla abundances. To illustrate the estimated differences in bacterial abundance when an associated clinical factor varies, the predicted percentage of sequencing reads (of the total in the bacterial community) have been calculated when a specified clinical factor is at its 25 % and 75 % quartile. Predicted values are indicated by bars, with whiskers indicating the 95 % confidence interval. Where multiple factors were found to influence an OTU (as indicated by the top bar), each factor is illustrated separately, with either then median or the most common categorical option being used as a base value for the additional clinical factors. The effects having mixed feeds (formula and breastmilk) are compared to breastmilk feeds alone.

### Term and premature infant faecal microbiota converge by two years of age

Whilst at six weeks of age the faecal microbiota of premature and term infants differs significantly, the microbial communities coverage towards a new structure by two years of age (Figure 2). Beta diversity distances between the premature and term communities are significantly reduced from the distances at six weeks (see Figure 3), although the distances between groups are still slightly higher than the intragroup distances. The communities no longer have significant differences in terms of community diversity (see Figure 4), with both term and premature infant faecal microbiota communities increasing significantly in diversity compared to six week samples. Only two low abundance OTUs were found to be associated with gestational age after correction for multiple tests (see Figure 8 and Supplementary Table 5). No association between phyla read abundance and gestational age was found (see Supplementary Table 6).

**Figure 8.**
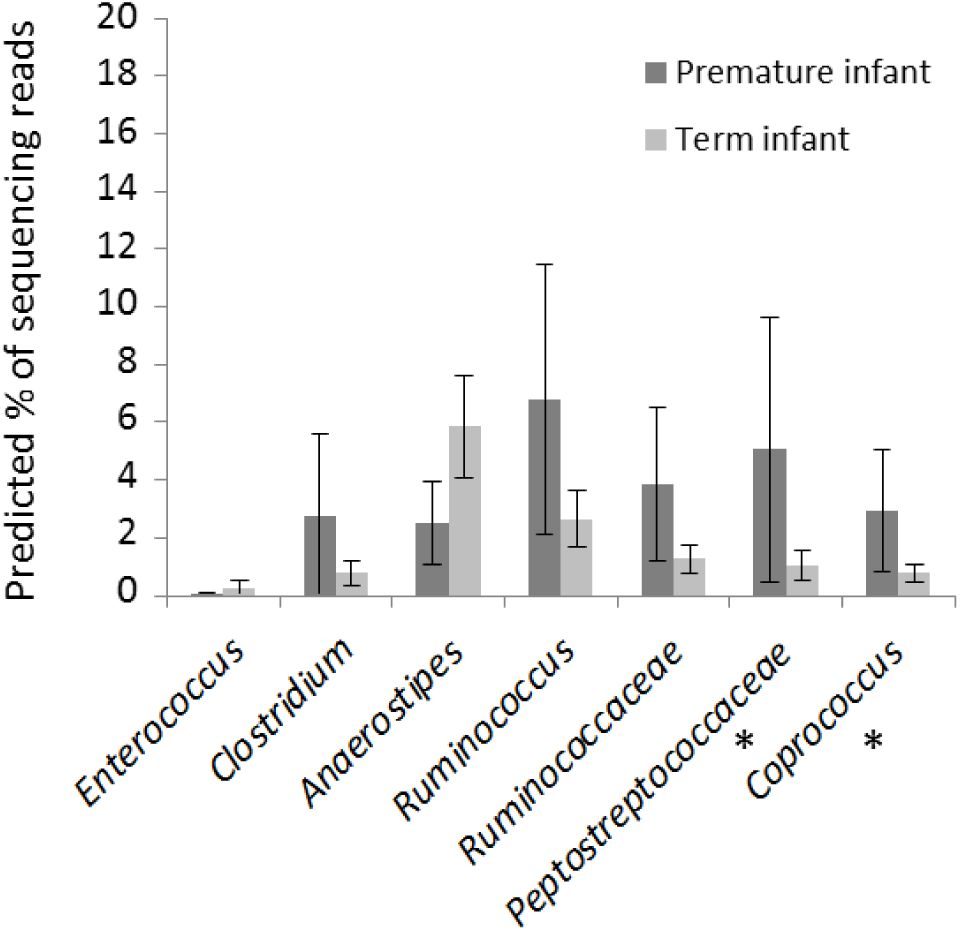
Bacterial OTUs that differ at two years of age in association with gestational age. The predicted percentage of sequencing reads (of the total in the bacterial community) for each OTU at two years of life have been calculated using the average gestational age of premature and term infants in this dataset (27.76 weeks and 40.35 weeks respectively). Predicted values are indicated by bars, with whiskers indicating the 95 % confidence interval. OTUs are displayed if significantly associated with age in GLMs. Stars indicate that this association remained significant after MHC.

While there was little evidence of significant differences between term and premature infants at two years of age, we investigated whether elements of the microbial community at two years of age was associated with clinical factors (as documented in Table 1; Birth demographics, Two year demographics and number of antibiotics courses by six months of age). OTUs and phyla that were significantly associated with clinical factors at two years of age are shown in Figure 9 (see Supplementary Tables 7 and 8).

**Figure 9.**
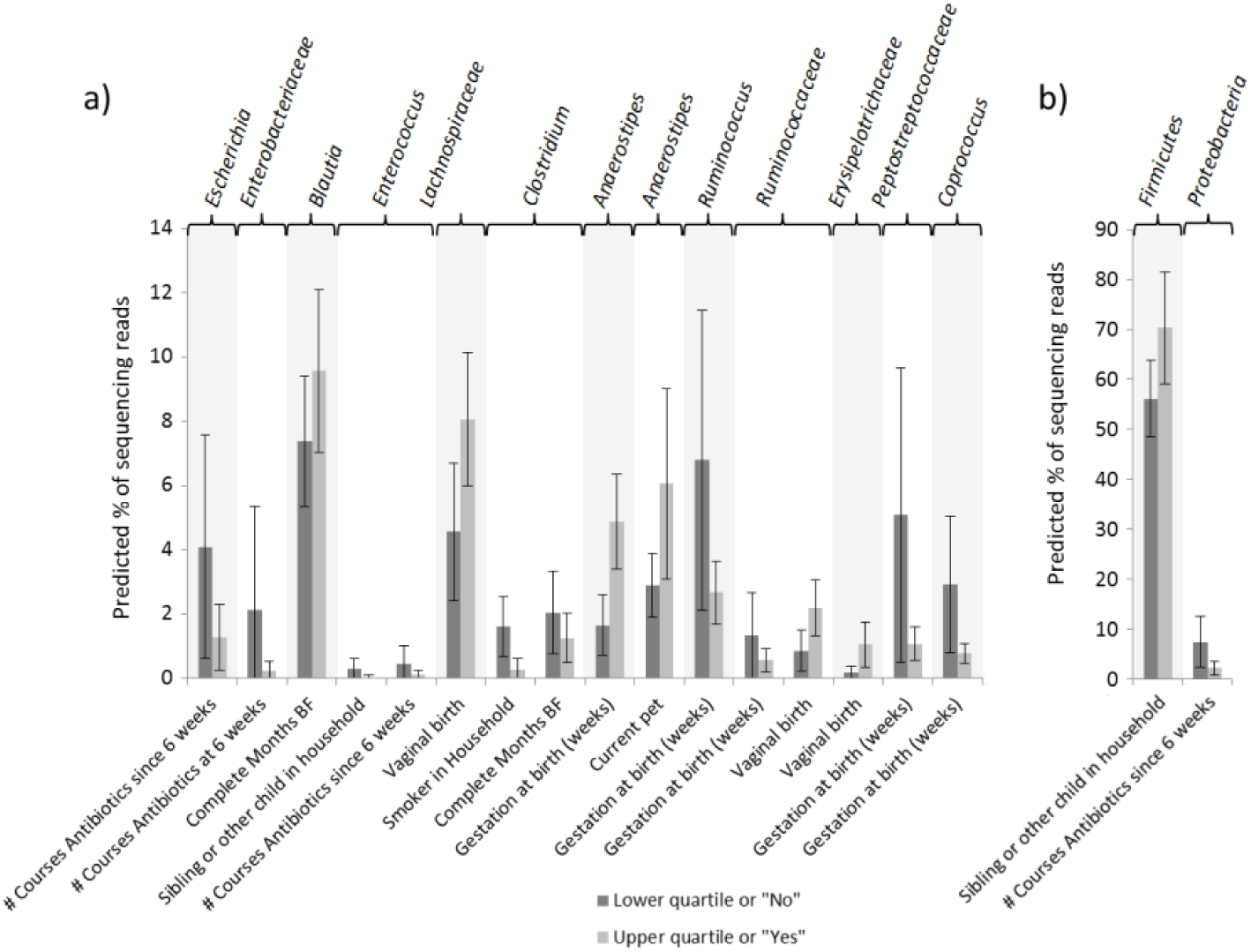
a) OTUs and their significantly associated clinical factors at two years of age. b) Phyla and their associated clinical factors at two years of age. Univariate and multivariate models were built where significant associations were found between clinical factors and OTU/phyla abundances. To illustrate the estimated differences in bacterial abundance when an associated clinical factor varies, the predicted percentage of sequencing reads (of the total in the bacterial community) have been calculated when a specified clinical factor is at its 25 % and 75 % quartile. Predicted values are indicated by bars, with whiskers indicating the 95 % confidence interval. Where multiple factors were found to influence an OTU (as indicated by the top bar), each factor is illustrated separately, with either then median or the most common categorical option being used as a base value for the additional clinical factors.

### Associations with early childhood conditions

Clinical data concerning the presence/absence of parent-reported wheezing and eczema by two years of age were collected for the enrolled infants. We sought to identify clinical factors and faecal bacterial signatures present both a six weeks and two years of life that were associated with the development of these two conditions. Univariate logistic regression models with a dependent variable of either wheeze (yes/no) or eczema (yes/no) were created for each set of bacterial signatures (OTUs and phyla) and clinical factors (as documented in Table 1; Birth demographics, Two year demographics and number of antibiotics courses by six months of age). Premature infants are known to have a lower incidence of eczema than term infants, hence gestation was included as a confounding factor in all univariate models.

At six weeks, no bacterial OTU or diversity measure was found to be associated with either wheezing or eczema by two years of age. At two years of age, increased Subdoligranulum reads were significantly associated with the development of wheeze (p = 0.0018, p = 0.0468 after MHC) although the OTU represents only a small proportion of sequencing reads from the microbial community (median 1%). Increased Firmicutes were also associated with both conditions (p = 0.002 for wheezing, p = 0.007 for eczema, p = 0.007 and p =0.029 respectively after MHC for four phyla tested). Significant associations were also found between increased weeks of birth gestation and eczema, and between increased courses of antibiotics since six weeks of life and the risk of wheeze (see Table 2). Factors significant to p = 0.2 after MHC were included in a multivariate models to predict the odds ratios of both wheezing and eczema for infants at the median and upper and lower quartiles of each variable.

The association between gestational age and the occurrence of eczema at two years is reflected in the lack of eczema cases reported in the premature infants in this study (8 cases in 37 infants, contrasted to 24 cases in the 37 term infants). We hypothesised that this relationship may mask other signals in the data, so repeated the analysis using only data from the 37 term infants. The positive association with Firmicutes and eczema remained (p = 0.035, odds ratio of 1.01 (95% CI 1.00, 1.01)) and *Faecalibacterium* was also found to be positively associated with eczema (p = 0.021, odds ratio of 1.03 (95% CI 1.01, 1.05)).

### Validation of factors associated with early childhood conditions

Given our findings that various microbial signals and clinical factors can be associated with development of eczema and wheeze, we sought to reproduce these results in a validation dataset drawn from 99 unmatched infant faecal samples at two years of age; 64 from infants born prematurely and 35 from infants born at term. ROC curves were created for the matched infant dataset (the discovery set) and the validation set (see Figure 10) using the variables retained in the multivariate models (see Table 2). For the occurrence of wheeze, an area under the curve (AUC) of 0.84 was obtained for the discovery set, although this was not accurately reproduced for the validation set (AUC = 0.63, IQR: 0.61-0.66, 1000 iterations). A similar observation was found for eczema, with the discovery set having an AUC of 0.79 and the validation set an AUC of 0.66 (IQR: 0.62-0.69, 1000 iterations).

**Figure 10.**
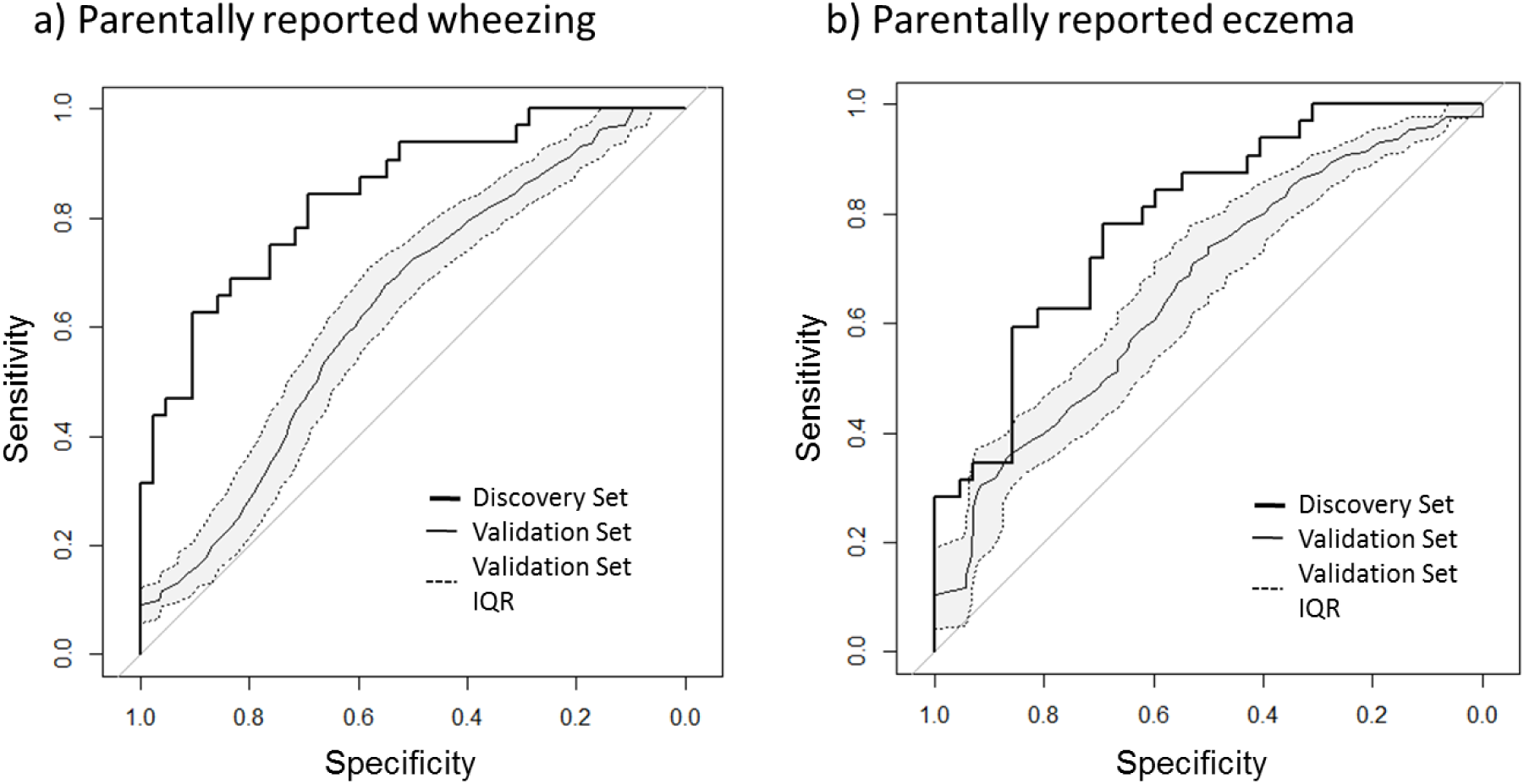
ROC curves illustrating the sensitivity and specificity of the factors identified through multivariate models. Factors identified for detecting a) parentally reported wheeze were Firmicutes reads, Subdoligranulum reads and the number of courses of antibiotics given since six weeks of age and b) parentally reported eczema were Firmicutes reads and gestation at birth. These factors were identified in the discovery set (37 matched premature and term infants), resulting in ROC curves shown in black. The validation set consisted of 30 term and 30 premature infants drawn randomly from a pool of 64 premature infants and 35 born at term 1,000 times. The solid grey lines indicate the median curves from these 1,000 iterations, and the dashed lines indicate the interquartile ranges.

## Discussion

### Convergence of premature and term infant gut microbiota by two years of age

Our data demonstrate the stark difference between the microbiota of the premature infant and the term infant gut microbiota at six weeks of age, with significant increases in *Bifidobacterium, Bacteroides* and *Lachnospiraceae* in in the term cohort and significantly increased *Enterobacteriaceae* and *Enterobacter* in the premature cohort. These differences are all absent at two years of age, indicating a convergence of all but a couple of low abundance bacterial OTUs. We similarly demonstrate a significantly higher microbial richness and diversity in term infants at six weeks compared to premature infants, with these differences absent by two years of age. The decreased microbial diversity of the premature gut during early months has been noted in other studies [8, 31]. Later studies have also demonstrated microbial convergence by five years of age [32], although our data suggest this process occurs even earlier in life. Whilst other studies have found difference in specific taxa up to four years of life [9], the relatively small numbers of premature infants observed are a limitation.

### Determinants of the gut microbiota at two years

Whilst gut microbiota was seen to converge across the broad categories of “term” and “premature” infants, clinical factors such as antibiotic use, gestational age at birth and delivery were still observed to be associated with differential abundances.

In comparison to other studies, [Fouhy 2019] similarly observed no associations between phyla at two years of age and gestational age at birth or mode of delivery. At a finer taxonomic resolution they did however observe increased streptococci to be associated with lower gestational age at birth, in line with our findings. Fouhy *et al* also noted an association between *Lachnospiraceae* and C-section deliveries at two years of age, in contrasts to our observations, although they noted the opposite in samples taken at four years of age. Our observed decrease in enterococci following neonatal antibiotic treatment has also been noted in other studies [Jia 2019], although we find evidence that this difference remains at two years of life. We also noted a significant increase in Proteobacteria at two years to be associated with increased antibiotic use; this may represent the delaying of the maturation of the microbiota following antibiotic treatment as suggested by Bokulich et al. 2017. Further follow-up of these infants would be required to confirm whether the microbiota will eventually fully converge of whether heavy antibiotic-use is associated with life-long increases in Proteobacteria, which could have health consequences given the association between these bacteria and inflammatory gut conditions such as IBD [Glassner 2020].

### Associations between wheeze and the gut microbiota at two years of age

We did not find an association between the microbiota at 6 weeks and later wheeze. This may be because of the inherent problems with parent-reported wheeze affecting our case classification and the early (2 years of age) diagnosis. As we follow the cohort to school age when a more rigorous assessment of respiratory function can be undertaken we may demonstrate an association. At 2 years we found a positive association between Firmicutes reads and *Subdoligranulum* and wheeze, and Firmicutes reads and eczema. However, the number of reads for *Subdoligranulum* is low overall and this result should be treated with some caution; although there is previous work in a small case control study demonstrating an association between sensitization to food allergens and *Subdoligranulum*[33].

### Associations with eczema at two years of age

For eczema, prospective studies have demonstrated in the majority reduced diversity, and a greater prevalence of clostridia species, and in individual studies an increase in *Escherichia coli*, or *Bacteroidetes* in early life correlating with eczema[34]. In our term infants we found a significant increase in *Faecalibacterium* in the faeces of cases of eczema. There are few cross-sectional studies at a similar age to our 2 year timepoint. Zheng et al[35] assessed the faecal microbiota at around one year in children with parent report of doctor-diagnosed eczema compared to controls; they did not find a difference in the phyla or diversity but found a significant difference in abundance of several genera with an increase in for example *Bifidobacterium, Streptococcus, Megasphaera* and *Haemophilus* in healthy infants, and an increase in for example *Veillonella, Lachnospiracaeae* and *Faecalibacterium* in infants with eczema. Another study, [36], (using microarray) has found a reduction in *Faecalibacterium prausnitzii* in atopic children, this apparent contradiction may be explained by dysbiosis as a subspecies level – Hong et al [37] found different subspecies were found in faeces from atopic dermatitis patients than that from controls. Nylund et al [38] in a microarray study assessing the effect of lactobacillus administration on eczema development found a higher diversity at 18 months of age in the faeces of children with eczema and a difference in abundance of taxa such as clostridium species.

We found an association between gestational age at birth and risk of parental report of eczema, such that babies born prematurely are less likely to have eczema. This has been reported elsewhere[39].

### Wheeze association with antibiotic courses since 6 weeks

Parental report of wheeze by 2 years was positively associated with the number of antibiotic courses since 6 weeks. There are a number of factors that could explain this association which has previously been found and explored further by others[40]. There may be recall bias such that parents of children who wheezed are more likely to report antibiotic use – cross-sectional studies have found a stronger association than longitudinal studies between wheeze and antibiotic use[41]. This may be relevant for the premature cohort however in the term cohort we visited infants regularly during the first two years and had corresponding GP notes so were able to reduce recall bias. Reverse causation may explain this association, as antibiotics may be prescribed for wheezing episodes rather than being the cause of such episodes. Other studies which have accounted for this factor have shown either no or a much smaller association between antibiotic use and wheeze[42], and in the term cohort overall accounting for antibiotics given wheeze the association became non-significant; we do not have sufficient information to conduct the same for the premature cohort but expect similar results. In addition, there may be an as yet unknown factor such as genetic predisposition to viral infections which increases the number of antibiotics prescribed and sequelae from early viral infections such as recurrent wheeze or asthma[43].

### Limitations

Our study relied on parental report of wheeze and eczema which is a significant limitation. Wheeze is a poorly understood term by parents which can be both underused[44] and overused[45], partly related to the first language used by parents and the age of the child. Even between medical professionals there can be discrepancy between defining a respiratory noise as wheeze[46]. Wheeze which is confirmed by a physician has been associated with higher specific airway resistance compared to that which is only reported by the parent.[47]

## Conclusions

Our data indicate a notable convergence of the premature and term infant gut microbiota by two years of age, yet some specific differences associated with specific clinical factors including delivery method, early antibiotic use and gestation at birth remain. These observations could be of great importance given the associations between the development of the gut microbiota and life-long health. The microbiota at six weeks of life was not found to be associated with wheeze at two years of age, whilst development of eczema was associated with early life gut microbial patterns. This study benefitted from comparably large numbers of infants and identical sample processing for the two cohorts.

## List of Abbreviations

AUC: area under the curve
CCA: Canonical Correspondence Analysis
CI: Confidence Interval
GLM: Generalised Linear Models
IBD: Inflammatory Bowel Disease
IQR: Interquartile Range
MHC: Multiple hypothesis corrections
NEC: Necrotising enterocolitis
OTU: Operational Taxonomic Unit

## Ethics approval and consent to participate

The study ‘Defining the Intestinal Microbiota in Premature Infants’ (ClinicalTrials.gov Identifier NCT01102738) was approved by West London Research Ethics Committee 2, United Kingdom (Reference number: 10/H0711/39). Parents gave written informed consent for their infant to participate in the study.

The study ‘Development Of Respiratory Microbiota In Children’ was approved by Riverside Ethics Committee, London, UK (Reference number: 12/LO/1362). Parents gave written informed consent for their infant to participate in the study.

## Consent for publication

Not applicable

## Data Availability

The dataset supporting the conclusions of this article is available in the European Nucleotide Archive repository, accession number PRJEB23362, https://www.ebi.ac.uk/ena/data/view/PRJEB23362.

## Authors’ contributions

KS and EP led the NeoM and DORMICe studies respectively. AS, KS, EP and EC performed DNA extractions and preparation for next-generation sequencing. AS performed the bioinformatics processing and statistical analyses. All authors contributed to the text and SK approved the final version.

## Acknowledgements

We thank all the participants and their families who enrolled in the NeoM and DORMICe and our colleagues at the Imperial College Healthcare National Health Service (NHS) Trust neonatal intensive care unit for supporting the study.

