## Supplementary Data for "Development of the gut microbiota during early life in premature and term infants"

| OTU Label | Coefficient (CI) | Exponentiated coefficient (CI) | P-value | Corrected P-value | Predicted % reads at 6 weeks of age and average gestation for premature infants (CI) | Predicted % reads at 6 weeks of age and average gestation for term infants (CI) |
|---|---|---|---|---|---|---|
| Bifidobacterium | 0.25 (0.16, 0.33) | **1.28** (1.18, 1.40) | <0.001 | <0.001 | **2.18** (0.44, 3.92) | **49.33** (15.53, 83.14) |
| Bacteroides | 0.34 (0.17, 0.47) | **1.40** (1.19, 1.60) | <0.001 | <0.001 | **0.20** (0.00, 0.44) | **13.93** (0.45, 27.40) |
| Enterobacteriaceae | -0.19 (-0.31, -0.09) | **0.83** (0.74, 0.92) | <0.001 | 0.002 | **24.87** (1.32, 48.42) | **2.23** (0.38, 4.07) |
| Enterococcus | -0.13 (-0.22, -0.04) | **0.88** (0.80, 0.96) | 0.002 | 0.051 | **13.06** (2.33, 23.79) | **2.54** (0.72, 4.37) |
| Lachnospiraceae | 0.34 (0.18, 0.49) | **1.40** (1.20, 1.63) | <0.001 | <0.001 | **0.05** (0.00, 0.14) | **3.80** (0.00, 7.96) |
| Enterobacter | -0.08 (-0.09, -0.07) | **0.92** (0.91, 0.92) | <0.001 | <0.001 | **8.88** (8.46, 9.30) | **3.06** (2.84, 3.27) |
| Staphylococcus | -0.08 (-0.17, 0.00) | **0.92** (0.84, 1.00) | 0.024 | 1.000 | **2.14** (0.57, 3.71) | **0.74** (0.26, 1.21) |

*Table 1 – Bacterial OTUs that are significantly association with gestational age at six weeks. Associations were sought using general linear models with a negative binomial distribution. Day of life of sampling was included in each model as a confounding factor. The exponentiated coefficient of indicates the proportional difference in an OTU for each additional week of gestational age at birth. The predicted percentage of sequencing reads (of the total in the bacterial community) for each OTU at six weeks of life have been calculated using the average gestational age of premature and term infants in this dataset (27.76 weeks and 40.35 weeks respectively). OTUs with a significant P-value prior to MHC are shown.*

| Phyla | Coefficient (CI) | Exponentiated coefficient (CI) | P-value | Corrected P-value | Predicted % reads at 6 weeks of age and average gestation for premature infants (CI) | Predicted % reads at 6 weeks of age and average gestation for term infants (CI) |
|---|---|---|---|---|---|---|
| Firmicutes | 0.00 (-0.04, 0.05) | 1.00 (0.96, 1.05) | 0.813 | 1 | **31.82** (18.75, 44.89) | **33.86** (21.76, 45.96) |
| Proteobacteria | -0.11 (-0.16, -0.06) | 0.90 (0.85, 0.94) | <0.001 | <0.001 | **60.92** (33.17, 88.66) | **15.37** (9.27, 21.47) |
| Bacteroidetes | 0.33 (0.17, 0.46) | 1.39 (1.19, 1.58) | <0.001 | <0.001 | **0.25** (0.00, 0.53) | **15.77** (1.23, 30.31) |
| Actinobacteria | 0.24 (0.16, 0.31) | 1.27 (1.17, 1.37) | <0.001 | <0.001 | **2.36** (0.68, 4.05) | **46.55** (18.06, 75.04) |

*Table 2 – Associations between bacterial phyla and gestational age at 6 weeks. Associations were sought using general linear models with a negative binomial distribution. Day of life of sampling was included in each model as a confounding factor. The exponentiated coefficient of indicates the proportional difference in an OTU for each additional week of gestational age at birth. The predicted percentage of sequencing reads (of the total in the bacterial community) for each OTU at six weeks of life have been calculated using the average gestational age of premature and term infants in this dataset (27.76 weeks and 40.35 weeks respectively).*

| OTU Label | Clinical Factor | Coefficient of factor (CI) | Exp(coeff) (CI) | P value | Clinical Factor value at: | | | Predicted % OTU reads when clinical factor is at: | | Value set for multivariate class (if applicable): |
|---|---|---|---|---|---|---|---|---|---|---|
| | | | | | Lower quartile | Median | Upper quartile | Lower quartile or "No" (CI) | Upper quartile or "Yes" (CI) | |
| Bifidobacterium | Number of antibiotics courses | -1.86 (-2.76, -0.94) | **0.16** (0.06, 0.39) | <0.001 | 0 | 1 | 1 | **35.81** (6.00, 65.63) | **5.59** (2.41, 8.77) | |
| Bacteroides | Feeds at sampling: mixed | 2.11 (-0.11, 6.79) | **8.26** (0.90, 884.68) | 0.007 | NA | NA | NA | **1.59** (0.22, 2.96) | **13.12** (0.00, 31.29) | 34.4 weeks gestation |
| Bacteroides | Gestation at birth (weeks) | 0.33 (0.19, 0.45) | **1.39** (1.20, 1.57) | <0.001 | 28.1 | 0 | 40.3 | **0.20** (0.00, 0.44) | **11.00** (0.00, 22.37) | Breast fed |
| Enterobacteriaceae | Feeds at sampling: mixed | 1.82 (0.48, 3.36) | **6.17** (1.62, 28.72) | 0.006 | NA | NA | NA | **3.12** (0.92, 5.33) | **19.28** (0.00, 41.59) | 34.4 weeks gestation |
| Enterobacteriaceae | Gestation at birth (weeks) | -0.26 (-0.38, -0.15) | **0.77** (0.69, 0.86) | <0.001 | 28.1 | 34.4 | 40.3 | **16.33** (1.41, 31.25) | **0.66** (0.08, 1.25) | Breast fed |
| Enterococcus | Number of antibiotics courses | 0.76 (0.06, 1.64) | **2.14** (1.06, 5.17) | 0.028 | 0 | 1 | 1 | **2.38** (0.40, 4.36) | **5.09** (2.27, 7.92) | |
| Lachnospiraceae | Number of antibiotics courses | -5.87 (-13.57, -0.97) | **0.003** (0.00, 0.38) | 0.001 | 0 | 1 | 1 | **7.00** (0.00, 19.89) | **0.02** (0.00, 0.07) | |
| Clostridium | Number of antibiotics courses | -1.55 (-2.57, -0.32) | **0.21** (0.08, 0.72) | 0.004 | 0 | 1 | 1 | **21.12** (0.00, 47.43) | **4.49** (0.78, 8.19) | |
| Enterobacter | Feeds at sampling: mixed | -6.19 (-7.96, -4.62) | **0.002** (0.00, 0.01) | <0.001 | NA | NA | NA | **29.43** (11.28, 47.57) | **0.06** (0.00. 0.15) | C-section, 34.4 weeks gestation |
| Enterobacter | Vaginal birth | -7.32 (-8.50, -6.33) | **0.001** (0.00, 0.002) | <0.001 | NA | NA | NA | **29.43** (11.28, 47.57) | **0.02** (0.00, 0.04) | Breast fed, 34.4 weeks gestation |
| Enterobacter | Gestation at birth (weeks) | -0.17 (-0.26, -0.08) | **0.85** (0.77, 0.92) | <0.001 | 28.1 | 34.4 | 40.3 | **84.50** (12.75, 100.00) | **10.96** (3.00, 18.91) | C-section & breast fed |
| Staphylococcus | Complete months breast feeding | -2.64 (-5.92, 0.22) | **0.07** (0.03, 1.25) | 0.014 | 0 | 1 | 1 | **14.95** (0.00, 46.45) | **1.07** (0.53, 1.61) | 34.4 weeks gestation |
| Staphylococcus | Gestation at birth (weeks) | -0.10 (-0.18, -0.02) | **0.91** (0.83, 0.98) | 0.009 | 28.1 | 34.4 | 40.3 | **1.96** (0.59, 3.33) | **0.60** (0.22, 0.99) | 1 month of breast feeding |

*Table 3 – OTUs and their associated clinical factors at six weeks. The coefficient of the association is shown with confidence intervals, which is exponentiated to give the proportional change (highlighted in bold) in OTU reads per unit of the clinical factor. To illustrate the effects of these shifts, predicted percentages of OTU read numbers have been calculated when the clinical factor is at its 25 % and 75*

*% quartile. Where multiple factors were found to influence an OTU the base values (either then median or the most common categorical option) for each clinical factor are given.*

| OTU Label | Clinical Factor | Coefficient of factor (CI) | Exp(coeff) (CI) | P value | Clinical Factor value at: | | | Predicted % OTU reads when clinical factor is at: | | Value set for multivariate class (if applicable): |
|---|---|---|---|---|---|---|---|---|---|---|
| | | | | | Lower quartile | Median | Upper quartile | Lower quartile or "No" (CI) | Upper quartile or "Yes" (CI) | |
| Proteobacteria | Gestation at birth (weeks) | -0.11 (-0.16, -0.06) | **0.90** (0.85,0.94) | <0.001 | 28.1 | 34.4 | 40.3 | **58.69** (32.61, 84.77) | **15.45** (9.34, 21.56) | |
| Bacteroidetes | Gestation at birth (weeks) | 0.32 (0.17, 0.44) | **1.38** (1.20, 1.55) | <0.001 | 28.1 | 34.4 | 40.3 | **0.24** (0.00, 0.51) | **12.19** (0.22, 24.170) | Breast fed |
| Bacteroidetes | Feed at sampling (mixed) | 1.97 (0.00, 3.87) | **7.16** (0.00, 368.38) | 0.009 | NA | NA | NA | **1.83** (0.34, 3.32) | **13.1** (0.00, 30.33) | 34.4 weeks gestation |
| Actinobacteria | # Courses Antibiotics | -1.74 (-2.52, -0.98) | **0.18** (0.08, 0.38) | <0.001 | 0 | 1 | 1 | **5.02** (0.04, 10.01) | **0.89** (0.05, 1.72) | C-section |
| Actinobacteria | Vaginal Delivery | 2.16 (1.05, 3.19) | **8.71** (2.87, 24.35) | <0.001 | NA | NA | NA | **0.89** (0.05, 1.72) | **7.72** (3.68, 11.76) | 1 course of ABX |

Table 4 – Phyla and their associated clinical factors at six weeks of age. *The coefficient of the association is shown with confidence intervals, which is exponentiated to give the proportional change (highlighted in bold) in OTU reads per unit of the clinical factor. To illustrate the effects of these shifts, predicted percentages of OTU read numbers have been calculated when the clinical factor is at its 25 % and 75 % quartile. Where multiple factors were found to influence an OTU the base values (either then median or the most common categorical option) for each clinical factor are given.*

| OTU Label | Coefficient (CI) | Exponentiated coefficient (CI) | P-value | Corrected P-value | Predicted % reads at 2 years of age and average gestation for premature infants (CI) | Predicted % reads at 2 years of age and average gestation for term infants (CI) |
|---|---|---|---|---|---|---|
| Enterococcus | 0.29 (0.05, 0.55) | **1.34** (1.05, 1.73) | 0.010 | 0.249 | **0.01** (0.00, 0.12) | **0.26** (0.00, 0.54) |
| Clostridium | -0.10 (-0.21, 0.01) | **0.90** (0.81, 0.99) | 0.023 | 1.000 | **2.75** (0.00, 5.57) | **0.78** (0.34, 1.21) |
| Anaerostipes | 0.07 (0.02, 0.12) | **1.07** (1.02, 1.13) | 0.006 | 0.157 | **2.50** (1.06, 3.94) | **5.87** (4.10, 7.64) |
| Ruminococcus | -0.08 (-0.14, -0.02) | **0.93** (0.87, 0.98) | 0.012 | 1.000 | **6.79** (2.11, 11.46) | **2.66** (1.67, 3.65) |
| Ruminococcaceae | -0.09 (-0.16, -0.03) | **0.91** (0.85, 0.97) | 0.003 | 0.082 | **3.84** (1.19, 6.49) | **1.27** (0.79, 1.75) |
| Peptostreptococcaceae | -0.13 (-0.22, -0.05) | **0.88** (0.80, 0.95) | 0.001 | 0.0367 | **5.08** (0.49, 9.66) | **1.06** (0.54, 1.58) |
| Coprococcus | -0.11 (-0.19, -0.04) | **0.90** (0.83, 0.96) | <0.001 | 0.0191 | **2.92** (0.80, 5.03) | **0.77** (0.46, 1.07) |

Table 5 – *Bacterial OTUs that are significantly association with gestational age at two years. Associations were sought using general linear models with a negative binomial distribution. Day of life of sampling was included in each model as a confounding factor. The exponentiated coefficient of indicates the proportional difference in an OTU for each additional week of gestational age at birth. The predicted percentage of sequencing reads (of the total in the bacterial community) for each OTU at two years of life have been calculated using the average gestational age of premature and term infants in this dataset (27.76 weeks and 40.35 weeks respectively). OTUs with a significant P-value prior to MHC are shown.*

| Phyla | Coefficient (CI) | Exponentiated coefficient (CI) | P-value | Corrected P-value | Predicted % of reads at 2 years of age and average gestation for premature infants (CI) | Predicted % of reads at 2 years of age and average gestation for term infants (CI) |
|---|---|---|---|---|---|---|
| Firmicutes | -0.00 (-0.02, 0.02) | **1.00** (0.98, 1.02) | 0.742 | 1 | **63.54** (48.77, 76.31) | **60.12** (53.23, 67.01) |
| Proteobacteria | 0.03 (-0.07, 0.11) | **1.03** (0.93, 1.11) | 0.583 | 1 | **2.89** (0.00, 6.05) | **4.01** (1.72, 6.29) |
| Bacteroidetes | -0.01 (-0.06, 0.03) | **0.99** (0.94, 1.03) | 0.574 | 1 | **29.33** (14.37, 44.29) | **25.09** (18.43, 31.75) |
| Actinobacteria | 0.02 (-0.02, 0.07) | **1.03** (0.98, 1.07) | 0.317 | 1 | **7.44** (3.15, 11.72) | **10.18** (7.14, 13.23) |

Table 6 – *Associations between bacterial phyla and gestational age at two years. Associations were sought using general linear models with a negative binomial distribution. Day of life of sampling was included in each model as a confounding factor. The exponentiated coefficient of indicates the proportional difference in an OTU for each additional week of gestational age at birth. The predicted percentage of sequencing reads (of the total in the bacterial community) for each OTU at two years of* *life have been calculated using the average gestational age of premature and term infants in this dataset (27.76 weeks and 40.35 weeks respectively).*

| OTU Label | Clinical Factor | Coefficient of factor (CI) | Exp(coeff) (CI) | P value | Clinical Factor value at: | | | Predicted % OTU reads when clinical factor is at: | | Value set for multivariate class (if applicable): |
|---|---|---|---|---|---|---|---|---|---|---|
| | | | | | Lower quartile | Median | Upper quartile | Lower quartile or "No" (CI) | Upper quartile or "Yes" (CI) | |
| Escherichia | # Courses Antibiotics since 6 weeks | -0.39 (-0.83, 0.03) | **0.68** (0.44, 1.03) | 0.001 | 0 | 1 | 3 | **4.09** (0.60, 7.59) | **1.26** (0.24, 2.28) | |
| Enterobacteriaceae | # Courses Antibiotics at 6 weeks | -2.25 (-3.90, -0.55) | **0.11** (0.02, 0.58) | 0.001 | 0 | 1 | 1 | **2.12** (0.00, 5.35) | **0.22** (0.00, 0.51) | |
| Blautia | Complete Months BF | 0.03 (0.01, 0.05) | **1.03** (1.01, 1.06) | 0.003 | 5 | 9 | 13 | **7.37** (5.34, 9.40) | **9.58** (7.04, 12.11) | |
| Enterococcus | Sibling or other child in household | -2.24 (-4.00, -0.58) | **0.11** (0.02, 0.56) | 0.017 | NA | NA | NA | **0.28** (0.00, 0.61) | **0.03** (0.00, 0.08) | 1 course of antibiotics since 6 weeks old |
| Enterococcus | # Courses Antibiotics since 6 weeks | -0.46 (-0.89, -0.06) | **0.63** (0.41, 0.94) | 0.023 | 0 | 1 | 3 | **0.45** (0.00, 1.02) | **0.11** (0.00, 0.26) | No other children |
| Lachnospiraceae | Vaginal birth (yes) | 0.57 (0.13, 0.98) | **1.77** (1.14, 2.68) | 0.009 | NA | NA | NA | **4.56** (2.42, 6.70) | **8.06** (5.99, 10.14) | |
| Clostridium | Smoker in Household | -1.79 (-3.04, -0.32) | **0.17** (0.05, 0.72) | 0.007 | NA | NA | NA | **1.60** (0.66, 2.53) | **0.27** (0.00, 0.61) | 9 months of breast feeding |
| Clostridium | Complete Months BF | -0.06 (-0.12, -0.00) | **0.94** (0.89, 1.00) | 0.023 | 5 | 9 | 13 | **2.03** (0.75, 3.32) | **1.25** (0.48, 2.02) | No smokers in the household |
| Anaerostipes | Gestation at birth (weeks) | 0.09 (0.04, 0.14) | **1.09** (1.04, 1.15) | <0.001 | 28.1 | 34.4 | 40.3 | **1.65** (0.70, 2.60) | **4.87** (3.40, 6.35) | No current pet |
| Anaerostipes | Current pet (yes) | 0.74 (0.26, 1.27) | **2.10** (1.29, 3.55) | 0.003 | NA | NA | NA | **2.89** (1.88, 3.89) | **6.06** (3.10, 9.02) | 34.4 weeks gestation |
| Ruminococcus | Gestation at birth (weeks) | -0.08 (-0.14, -0.02) | **0.93** (0.87, 0.98) | 0.012 | 28.1 | 34.4 | 40.3 | **6.79** (2.11, 11.46) | **2.66** (1.67, 3.65) | |
| Ruminococcaceae | Gestation at birth (weeks) | -0.07 (-0.14, -0.01) | **0.93** (0.87, 0.99) | 0.006 | 28.1 | 34.4 | 40.3 | **1.33** (0.01, 2.66) | **0.56** (0.19, 0.93) | C-section |
| Ruminococcaceae | Vaginal birth (yes) | 0.94 (0.22, 1.62) | **2.56** (1.24, 5.03) | 0.025 | NA | NA | NA | **0.85** (0.21, 1.49) | **2.18** (1.31, 3.05) | 34.4 weeks gestation |
| Erysipelotrichaceae | Vaginal birth (yes) | 1.85 (0.56, 3.01) | **6.37** (1.76, 20.35) | 0.002 | NA | NA | NA | **0.16** (0.00, 0.38) | **1.04** (0.34, 1.74) | |
| Peptostreptococcaceae | Gestation at birth (weeks) | -0.13 (-0.22, -0.05) | **0.88** (0.80, 0.95) | 0.001 | 28.1 | 34.4 | 40.3 | **5.08** (0.49, 9.66) | **1.06** (0.54, 1.58) | |
| Coprococcus | Gestation at birth (weeks) | -0.11 (-0.19, -0.04) | **0.90** (0.83, 0.96) | <0.001 | 28.1 | 34.4 | 40.3 | **2.92** (0.80, 5.03) | **0.77** (0.46, 1.07) | |

*Table 7 - OTUs and their associated clinical factors at two years. The coefficient of the association is shown with confidence intervals, which is exponentiated to give the proportional change (highlighted in bold) in OTU reads per unit of the clinical factor. To illustrate the effects of these shifts, predicted percentages of OTU read numbers have been calculated when the clinical factor is at its 25 % and 75 % quartile. Where multiple factors were found to influence an OTU the base values (either then median or the most common categorical option) for each clinical factor are given.*

| OTU Label | Clinical Factor | Coefficient of factor (CI) | Exp(coeff) (CI) | P value | Clinical Factor value at: | | | Predicted % OTU reads when clinical factor is at: | |
|---|---|---|---|---|---|---|---|---|---|
| | | | | | Lower quartile | Median | Upper quartile | Lower quartile or "No" | Upper quartile or "Yes" |
| Firmicutes | Sibling or other child in household | 0.23 (0.06, 0.39) | **1.25** (1.06, 1.48) | 0.007 | NA | NA | NA | **56.12** (48.50, 63.75) | **70.32** (59.19, 81.44) |
| Proteobacteria | # Courses Antibiotics since 6 weeks | -0.41 (-0.69, -0.11) | **0.67** (0.50, 0.89) | <0.001 | 0 | 1 | 3 | **7.35** (2.24, 12.46) | **2.16** (0.74, 3.58) |

*Table 8 - Phyla and their associated clinical factors at two years of age. The coefficient of the association is shown with confidence intervals, which is exponentiated to give the proportional change (highlighted in bold) in OTU reads per unit of the clinical factor. To illustrate the effects of these shifts, predicted percentages of OTU read numbers have been calculated when the clinical factor is at its 25 % and 75 % quartile. Where multiple factors were found to influence an OTU the base values (either then median or the most common categorical option) for each clinical factor are given.*
